# Frequency specific network effective connectivity: ERP analysis of recognition memory process by directed connectivity estimators

**DOI:** 10.1101/739573

**Authors:** Mohammad Javad Darvishi Bayazi, Ali Motie Nasrabadi, Tim Curran

## Abstract

Various processes occur in memory retrieval in recognition memory and it is necessary to investigate memory brain function. Most of the research in past decades have focused on particular brain region function, but the interaction between these has a major role in human cognition. In this study, we used the memory retrieval task to investigate the underlying mechanism of recognition memory. The connectivity between brain regions is estimated from scalp electroencephalography signals that were recorded from twenty-three healthy subject participated in recognition memory task to correctly classify old/new words. Multivariate autoregressive models (MVAR) are used for the determination of Granger causality to estimate the effective connectivity in the time-frequency domain. We use GPDC and dDTF methods because they have almost resolved the previous problems in estimations. Results show that brain regions in the old condition have greater global connectivity in the theta and gamma band compared to the new words retrieval. Connectivity within and between the brain’s hemisphere may be related to correct rejection. The left frontal has a crucial role in recollection. theta and gamma specific connectivity pattern between temporal, parietal and frontal cortex may disclose the retrieval mechanism. old/new comparison resulted in the different patterns of network connection. These results and other evidence emphasize the role of frequency of causal network interactions in the memory process.

## 1 Introduction

The next move in the advancement of neuroscience is creating neuro-cognitive models which can describe the brain region’s dynamics and interaction pattern on a macroscopic scale. Recent progress in cognitive neuroscience has concentrated on the part of inter-areal interaction between different specialized brain regions and practical interactions in human cognition [51, 62, 47]. Since many studies on brain memory have reported several brain regions and frequency bands, it is necessary to investigate the pattern of activation of these regions in frequency bands.

Memory is a capacity of mind that can be encoded with the old/new information. Recognition memory test aims to recognize if a stimulus was previously presented individually. It was worthwhile that cognitive neuroscientists had shown more interest in over the past decades. In our work, Event-related brain potentials (ERPs) are used to separate two different events as old/new words [16].

Episodic memory formation is a complex neurocognitive process that is associated with large-scale neuronal activity distributed across the cortex. However, neuronal mechanisms based on the effective connectivity underlying the coordination of this anatomically distributed processing have remained largely unknown. The studies done on neuroimaging and patient injury for several decades showed a connection between the roles of particular regions to episodic memory retrieval like the prefrontal cortex [12, 20], the medial temporal lobe [21, 41, 49] and some parietal cortex sub-regions [33, 50, 56]. Moreover, the activities across brain regions, whether distributed or coordinated, are considered Important to the memory retrieval processes [14, 22, 45]. coincident activity in the local field potential (LFP) is related to coordinating the process as well [25].

Following the previous achievements, frequency multiplexing of brain regions involved in episodic memory retrieval has also been studied recently [58], but effective connectivity has been less regarded [38, 52, 17].

In addition, Granger introduced causality by the mathematical definition of time series xi(t), which is Granger-cause xj(t) if the past knowledge of xj(t) improves prediction of xi(n) significantly [28], although in [29] he mentioned that this casualty estimation is valid when there is no other time series influence on the process, to address this issue Multivariate autoregressive model (MVAR) can be assumed as Granger causality principle. Several measures, such as Granger Causality Index (GCI), Directed Transfer Function (DTF), Partial Directed Coherence (PDC) and their modifications are defined in the context of Multivariate Autoregressive Model and based on Granger causality principle [11].

The linear connection between two-time series can be estimated by PDC and DTF as it can disclose the crucial aspects of functional connectivity in neuroscience due to the central role played by neural rhythms (*δ, θ, α, β, γ*) that are of paramount physiologic relevance [7].

Functional connectivity measures the statistical connectivity between neural sources directly from the brain activity data [23]. PDC and DTF determine Granger Causality that is defined based on the multiple time-series “directed coherence (DC)” [50] and in multichannel data when more pairs than those of time series are simultaneously analyzed. One the other hand, the generalization of DC was introduced for multiple time series [34] (see [1, 4] for a comparison). However, the advantage of PDC and directed direct transform function (dDTF) are cascade connection estimation from direct connection due to the notion of partial coherence. For example, where node 1 links to node 2 and node 2 is connected to node 3, nodes 1 and 3 are not directly connected and PDC does not show the connection between the nodes 1 and 3, but evidence shows some other methods cannot do this [4].

Several works distinguish that memory process is in low-frequency bands [36, 32] Memory-related connectivity in measurement signals can be estimated based on PDC as well. Formerly, Generalized Partial Directed Coherence (GPDC) was estimated by using multivariate auto-regressive (MVAR) models. In this work, we focused on memory recognition task, using GPDC and dDTF methods of effective connectivity measurement.

Here in this paper, we are primarily concerned with the information flow network of the human brain during episodic memory retrieval. We estimate effective connectivity old/new memory task.

## 2 Material and Method

### 2.1 Database

We used a dataset which twenty-three of Colorado University students have participated in the recognition memory task, and it is a Combined Pharmacological and Electrophysiological study recording by [16]. All right-handed subjects were native English speakers and in the weigh group of 83 kg. A lot of considerations have been met, for more detail see [16].

The number of the word stimuli was 240 low-frequency English that two classes in each condition (old/new) assumed, which included 120-word sets randomly. In the Center of a computer monitor displayed each word for 4 s, with a one-second interval from inter-word. The subjects studied 120 words, followed by 70 min of interpolated activity including several cognitive tasks and Sensor Net setup (MN, 70.8 min), and also by the 240 word recognition memory test. Each trial began with a randomly determined 500–1000 ms fixation point, followed by a test-word presentation for 2000 ms. The subjects were instructed to withhold their response until a question mark appeared immediately after the test-word offset. The responses were delayed in this manner to minimize the response-related ERP effects. The responses were made by pressing keys on a response box with the first finger of each hand. The assignment of hands to old/new response was counterbalanced across subjects. A 500 ms inter trial interval followed each response.

EEG was recorded from 128-channels Geodesic Sensor Net by high-input impedance amplifier during the recognition memory task. The recorded scalp voltage was Amplified analogically and filtered by 0.1–g100 Hz band-pass and sampled at 250 Hz.

### 2.2 Data Pre-processing

The data were processed by EEGLAB toolbox [18] and custom-written codes in MATLAB. In the first step the artifact was detected and rejected by eyes, then the DC offset of signals were calculated, and the baseline of all channels was subtracted. Many analysts automatically perform a notch filter at 60 Hz to remove line noise. Such notch filters often use a notch width of 10 Hz or larger, resulting in significant signal distortion in the frequencies between 50 and 70 Hz. This distortion should not be an issue for analyses that immediately apply a low-pass filter to the signal, say at 40 Hz, but may preclude certain high-frequency studies. A number of studies have also revealed that non-casual low-pass filtering is able to change ERP onsets considerably [6, 10, 26].

Moreover, [6] it has been shown that filtering, especially low-pass one, can be of a damaging effect on Granger and other connectivity calculation. Afterwards so as to prevent it, a multi-taper decomposition method to detect and remove line noise constituents when minimizing background signal distortion was applied leverage regular ways from the CLEANLINE EEGLAB plug-in based on [42] was used.

There is also concern about the impact of high-pass filtering on ERP and connectivity analysis. The concern and a number of essays about the danger of filtering for ERP processing [48, 55, 53, 61] caused not to use high pass filtering. The filtering process is not used to remove the artifacts from the EEG signals and thus other processing is needed. So, such the high-variance artifacts in the signals as eye movement, blinking, muscle activity, and environmental noise, were removed by applying artifact subspace reconstruction (ASR) [44] which have been provided in the EEGLAB toolbox. In this way, the ASR threshold was set at 5 times of standard deviation. The ASR approach consists of a sliding window Principal Component Analysis (PCA), which statistically interpolates any high variance signal components greater than threshold referring to the covariance of an automatically detected data section that is relatively free of artifacts. The bad channel removed and then interpolated by spherical interpolation.

The data were re-referenced to the average of channels. To prevent the bad effect of noisy channels, we were cleaned all bad ones in previous steps.The epochs were extracted from 100 ms before to 1,000 ms after stimulation. The baseline of ERPs was estimated from 100 ms to Stimulation onset, and then subtracted from the whole trial.

We select eight regions of interest [Left, Anterior, Inferior (LAI) and Right, Anterior, Inferior (RAI) and Left, Anterior, Superior (LAS) and Right, Anterior, Superior (RAS) and Left, Posterior, Inferior (LPI) and Right, Posterior, Inferior (RPI) and Left, Posterior, Superior (LPS) and Right, Posterior, Superior (RPS)] Figure(1) from data channels according that suggested in [16, 59, 63].

**Figure 1:**
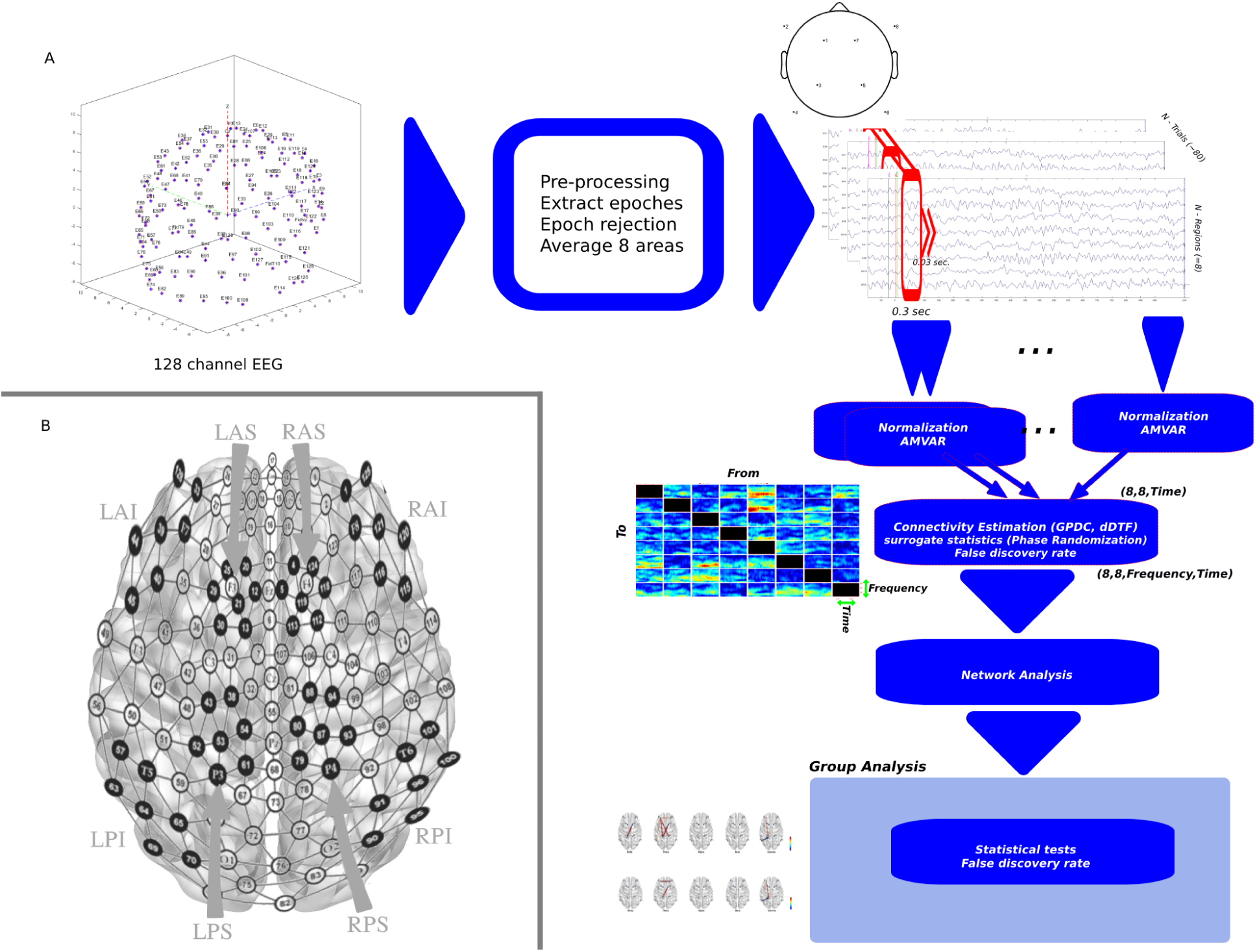
A) Schematic of overall method consist of pre-processing, sliding-window AMVAR modeling, connectivity estimation and validation and network analysis. B) Co-registration between Geodesic Sensor Net. and 10-20 system, the arrows mention eight regions of interest [Left, Anterior, Inferior (LAI) and Right, Anterior, Inferior (RAI) and Left, Anterior, Superior (LAS) and Right, Anterior, Superior (RAS) and Left, Posterior, Inferior (LPI) and Right, Posterior, Inferior (RPI) and Left, Posterior, Superior (LPS) and Right, Posterior, Superior (RPS)]

Afterward, we fit an adaptive multivariate auto-regressive (AMVAR) model and estimate connectivity. The mathematical explanation is as shown in 2.3 2.4.

A segmentation-based AMVAR applies windowing techniques. First, it extracts the parts of multi-channel data-set by sliding the window of which the length W, and fits VAR[p] model to these data (as seen in figure 1).

The Estimation of windows’ optimal length is a key choice in this method to have well temporal smoothing, locally stationary, sufficient amount of data in each window, processing dynamics, and Neuro-physiology. Short windows can improve the locally stationary of the data [6]. We apply window with 0.3 second length for MVAR model.

To validate our fitted model, we have to check the whiteness of the residuals, percentage consistency, and model stability for each (or a random subset) of our windows. Residual whiteness tests include portmanteau and auto-correlation tests to be correlated in the residuals of the model. Here, we benefit Ljung-Box, Box-Pierce, and Li-McLeod multivariate portmanteau tests and a simple auto-correlation function test.

For connectivity estimation, we used GPDC and dDTF estimator. These work with the phase difference between signals; only when there is a phase difference between signals, they have a non-zero value. Volume conduction affects the magnitude of the electrodes and has a zero-phase propagation; therefore, volume conduction does not generate a phase difference between channels. For this reason, theoretically, connectivity results should not be affected by volume conduction. Practically, it has some influence, e.g. increasing the noise level, however, this influence is not critical [35].

### 2.3 Generalized Partial Directed Coherence

Partial Directed Coherence is known as a linear quantifier which is connected with the multivariate relationship between observed time series in the frequency domain at the same time, to apply in functional connectivity inference in neuroscience. GPDC stands for the PDC re-definition, which improve PDC’s estimations under outlines with extremely unbalanced and predictive modeling errors (additive Gaussian noise). The GPDC is stronger versus the scale of time series [3]. If x(n) is assumed to be a matrix of N simultaneously observed time series as below:

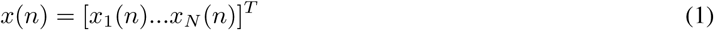

It can be modeled by a Multivariate Auto-regressive Model which is defined as:

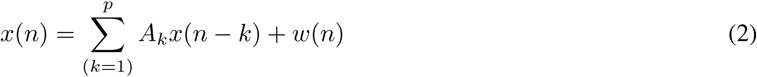

Where *p* is model order, *A_k_* is the matrix of coefficients *α_i_j* (*k*) standing how much the time series sample of *j^th^* affects *ith* time series at lag *k, w*(*n*) is also given as:

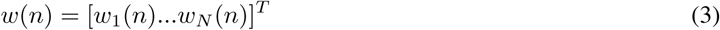

In fact, *w*(*n*) Is the vector of model’s additive Gaussian noise vector with zero mean and covariance matrix Σ *w*. The PDC is defined as:

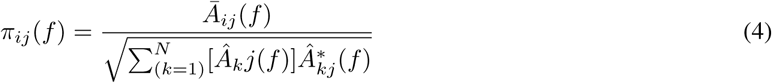

where f is the normalized frequency in the interval [-0.5, 0.5] where:

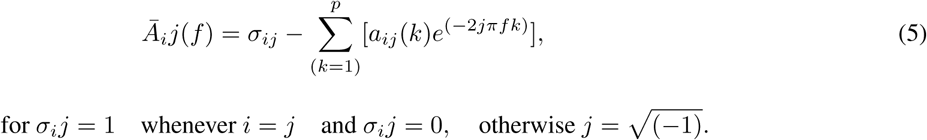

In [30], Baccala et al. Clearly showed a numerical problem in PDC with several examples. As one of the time series is scaled, PDC cannot estimate either. This necessarily results in the use of weighing the functions and the generalized definition of PDC. To settle the numerical issue, a new partial directed coherence estimator was defined as below:

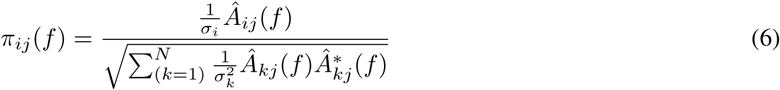

consequently, it follows as below:

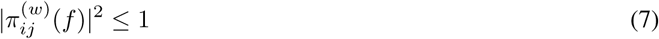

And

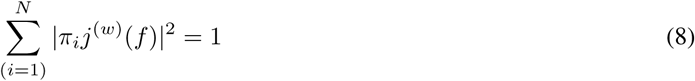

In the GPDC definition the normalization (7, 8) is the same as the PDC definition (4) (see [4]).

L. Astolfi et al. applied this method to Predefined patterns of cortical connectivity and simulated it, where high-resolution EEG Data were recorded during the well-known strop paradigm and therefore, the results indicated that the method correctly estimates connectivity patterns under reasonable conditions [1].

### 2.4 direct Directed Transfer Function (dDTF)

A multivariate time series like x(n) in equitation (1) could model with equitation (2) also by changing *k* = 0 and *A_K_* the model is:

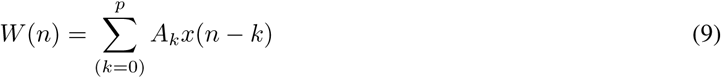

By applying z transform:

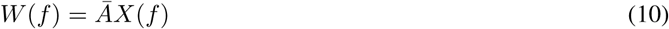

So:

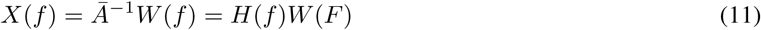

Partial coherence define as:

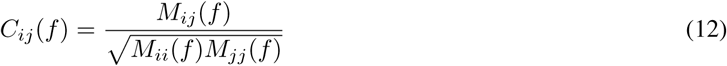

where *M_ij_* is a minor of spectral matrix. The dDTF [37] estimate direct influence between time series is defined as:

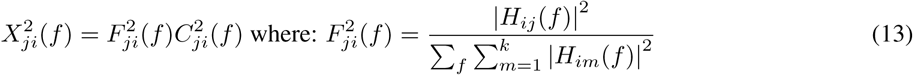

### 2.5 Statistical Analysis

So as to evaluate the significance of our connectivity results, we employed statistical hypothesis testing for each individual pair-wise connection within a multi-channel EEG dataset using a null distribution that we generated from the signal itself. The Statistical routines in this work fall into two categories, within-subject tests and group-level tests. Within-subject statistical test examines whether the expected value of the measure is significantly greater than zero (Hnull), here the phase randomization method was used. Group-level test, testing whether expected values of the measure differ under two experimental conditions (HAB), here the t-student test was applied. Phase randomization [54] is a method for estimating the empirical probability distribution of an estimator under a null hypothesis of no information transfer, temporal correlation, or phase synchrony, while striving to preserve spectral amplitude and other properties of the system. The procedure repeatedly randomizes the phases of all time-series, preserving the amplitude distribution. For each phase-randomized ensemble, re-compute GPDC estimator. Repeating this procedure many times produces the desired null distribution. It is implemented in SIFT [19].

The frequency-domain information flow estimators rely on the relative phases of the multivariate time-series, so any non-zero information flow in the surrogate data-set is because of chance, and the value of the estimator is compared to the null distribution quantities to obtain a probability estimate. Virtually, adjusting the significance thresholds to account for multiple comparisons is important so a basic multiple comparison correction by the False Discovery Rate (FDR) method [8], Bonferroni’s method was applied.

After calculating the valid connectivity in individuals, group-level test (T student test (p < 0.05)) was done between each Feature pair. In this step we apply paired t-test to every edge in six frequency band (Delta [0 4], Theta [4 8], Alpha [8 13], Beta [13 30], Gamma-Low [30 55] and Gamma-High [65 100]). To address the The problem of multiple comparisons [39] we used MATLAB implementation of false discovery rate method [9] with FDR=0.20.

### 2.6 Network Analysis

For network analysis, we investigate several aspects of the networks that estimated with the GPDC and dDTF. The time-frequency weighted networks are averaged across time and then averaged across every frequency band. The averaged networks are shown for both old and new conditions. Also, the significant edges between old/new are extracted and the significant difference (*connectivity_o1d_* — *connectivity_new_*) is shown, this difference is normalized for visualization, the color of edges present this difference and colors limited between blue (−1) and red (+1) so red color means the old condition have stronger connection than new and blue shows new have stronger connection at that edge. To evaluate node importance, the inflow/outflow of nodes calculated, the below formula is used to estimate inflow/outflow features:

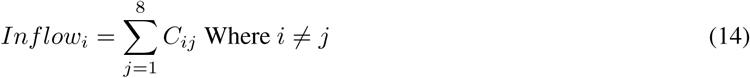

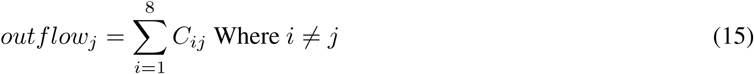

Where *C_ij_* is the strength of the connection that estimated by GPDC/dDTF methods, *C_ij_* shows strength of connection from node *j* to node *i*. Then the significant difference between old/new is shown.

## 3 Results

After pre-processing, the data were fitted to time-varying adaptive multivariate auto-regressive (TV-AMVAR) model and then the models were validated. Then the connectivity between eight regions was estimated by GPDC in the time-frequency domain, afterward, the non-valid (non-significant in Within-subject statistical test) was cleaned by using the phase randomization test. In the end, then group-level statistical analysis was performed for every edge between the eight region and the significant edges is presented in the Figure (2) for the GPDC method and dDTF results are shown in Figure (3). To visualize the average connectivity within the condition are calculated and the difference between these quantities between old and new are plotted by colors of edges as the red one means the old have a stronger connection while blue arrow shows the new is stronger than old, figures show in every frequency band across all time of trial.

**Figure 2:**
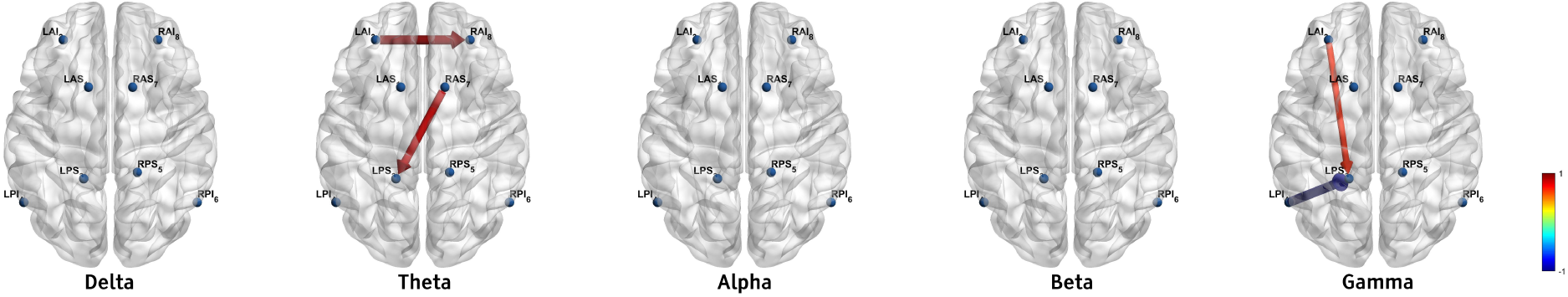
Significant difference between old/new by using the GPDC estimator, (*connectivity_old_* — *connectivity_new_*), this difference is normalized for visualization, the color of edges present this difference and colors limited between blue (−1) and red (+1) so red color means the old condition have stronger connection than new and blue shows new have stronger connection at that edge, From left to right brain maps show frequency band delta, theta, alpha, beta and gamma respectively.

**Figure 3:**
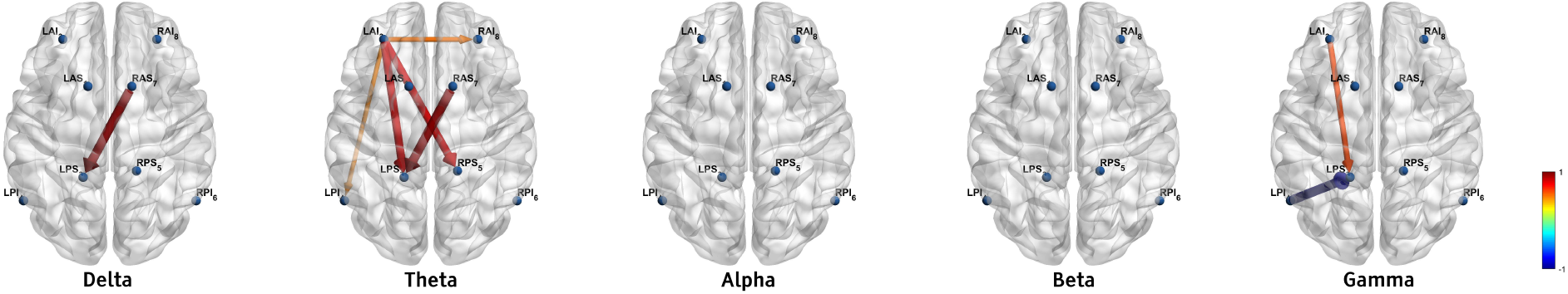
Significant difference between old/new by using the dDTF estimator, (*connectivity_old_* — *connectivity_new_*), this difference is normalized for visualization, the color of edges present this difference and colors limited between blue (−1) and red (+1) so red color means the old condition have stronger connection than new and blue shows new have stronger connection at that edge, From left to right brain maps show frequency band delta, theta, alpha, beta and gamma respectively.

### 3.1 Network Edges

We visualize regions connectivity in frequency bands. For better visualization, we only show the connections that have significant differences between the two old/new states. In Figure (2) the strength of connections is specified by the color of the arrows.

In the delta band, it is observed that no connection is significant between two old/new condition. In the theta band, the connection from LAI to RAI and RAS to LPS are stronger in the old state. But in the Alpha and beta as delta band, No significant difference was found. In the gamma band, it is observed that the connection from LAI to LPS is stronger in the old state and the forward connection from LPI to LPS is stronger in the new words.

The significant differences between old/new states are in the theta and gamma band. In the theta band, we observe Inter-hemispheric and in the gamma band intra-hemispheric Connectivity. Most of the connections in old condition are stronger than new except the connection from LPI to LPS in the gamma band.

For the dDTF as another estimator, We visualize regions connectivity in frequency bands. For better visualization, we only show the connections that have significant differences between the two old/new states. In Figure (3) the strength of connections is specified by the color of the arrows. The results are almost the same as the GPDC.

In the delta band, it is observed that one significant connection between two old/new condition is from RAS to LPS.In the theta band, The LAI node shows a very important role in our memory task, the figure shows the connection from LAI to RAI, RPS, LPS, LPI. Also, we observe connection from RAS to LPS, all connections in this band are stronger in the old state.In the Alpha and beta bands, No significant difference was found. In the gamma band, it is observed that the connection from LAI to LPS is stronger in the old state and the forward connection from LPI to LPS is stronger in the new words.

As we observe, The most significant differences between old/new states are in The theta an gamma band. In the theta band, we observe inter and Intra-hemispheric and in the gamma band intra-hemispheric Connectivity. Most of the connections in old condition are stronger than new except the connection from interior to superior of the left posterior region in the gamma band.

### 3.2 Channel Inflows and Outflows

After estimating the network of each subject and performing a statistical test to eliminate chance connection, we calculate the inflow/outflow of each channel and compare them in two condition, between group old /new states. In this way, we summed all outflow from each channel to other channels over all times and frequency bands as outflow and summed all inflow for each channel from other channels over time and frequency band as inflow. The GPDC and dDTF show some significant difference between the new/old states in the delta, theta, alpha and gamma band.

Regarding the equation (7, 8) and evidence that reported in [57], for the GPDC, the results of output are more valuable. Also, because the dDTF values are normalized to the inflow of channel so the outflow of dDTF is more valuable.

The GPDC inflow in delta is significantly higher in RAI and LPS in old words, and it is significant in RAI at theta frequency band. Also, the outflow is higher in anterior inferior regions in theta and alpha and is higher in gamma at LPS and RPI but lower in LPI.

The dDTF outflow in theta is significant and have higher quantity in LAI in old words. Also, the inflow is higher in LPS, LPI, RAI and LAI regions in delta, theta and alpha bands.

### 3.3 Networks in old/new states

In the section (3.1) we just show the significant difference between two conditions (Figure (6, 7)), in this section we show the average network of old/new and their different. this results show the overall network for old/new is very similar and the differences are in week connections.

**Figure 4:**
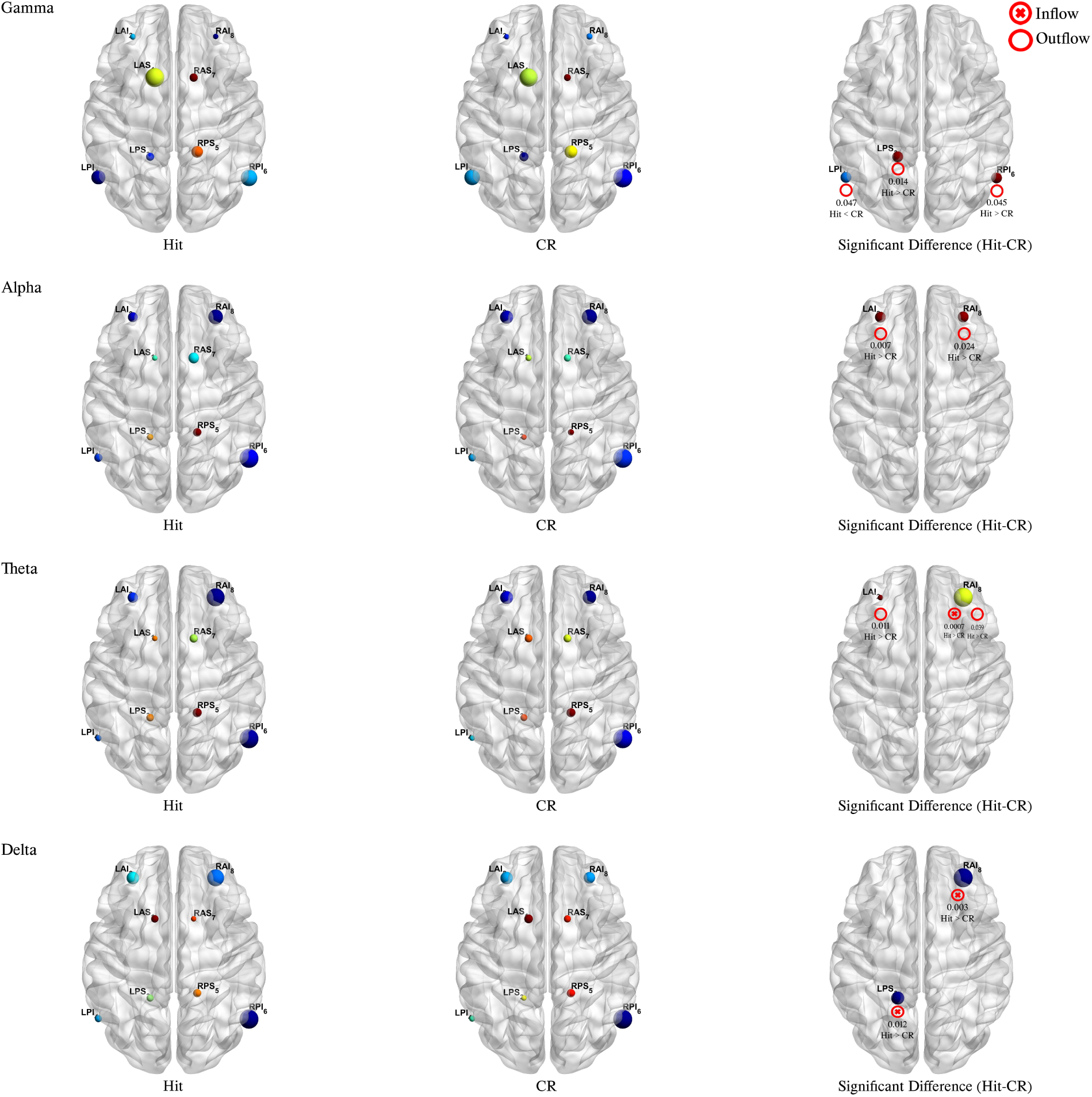
The information flow of regions estimated by GPDC in old/new (as Hit/CR) and the significant nodes between two conditions. The color of nodes present the outflow intensity and the node size shows inflow. Also the significant difference showed by special signs for inflow/outflow.

**Figure 5:**
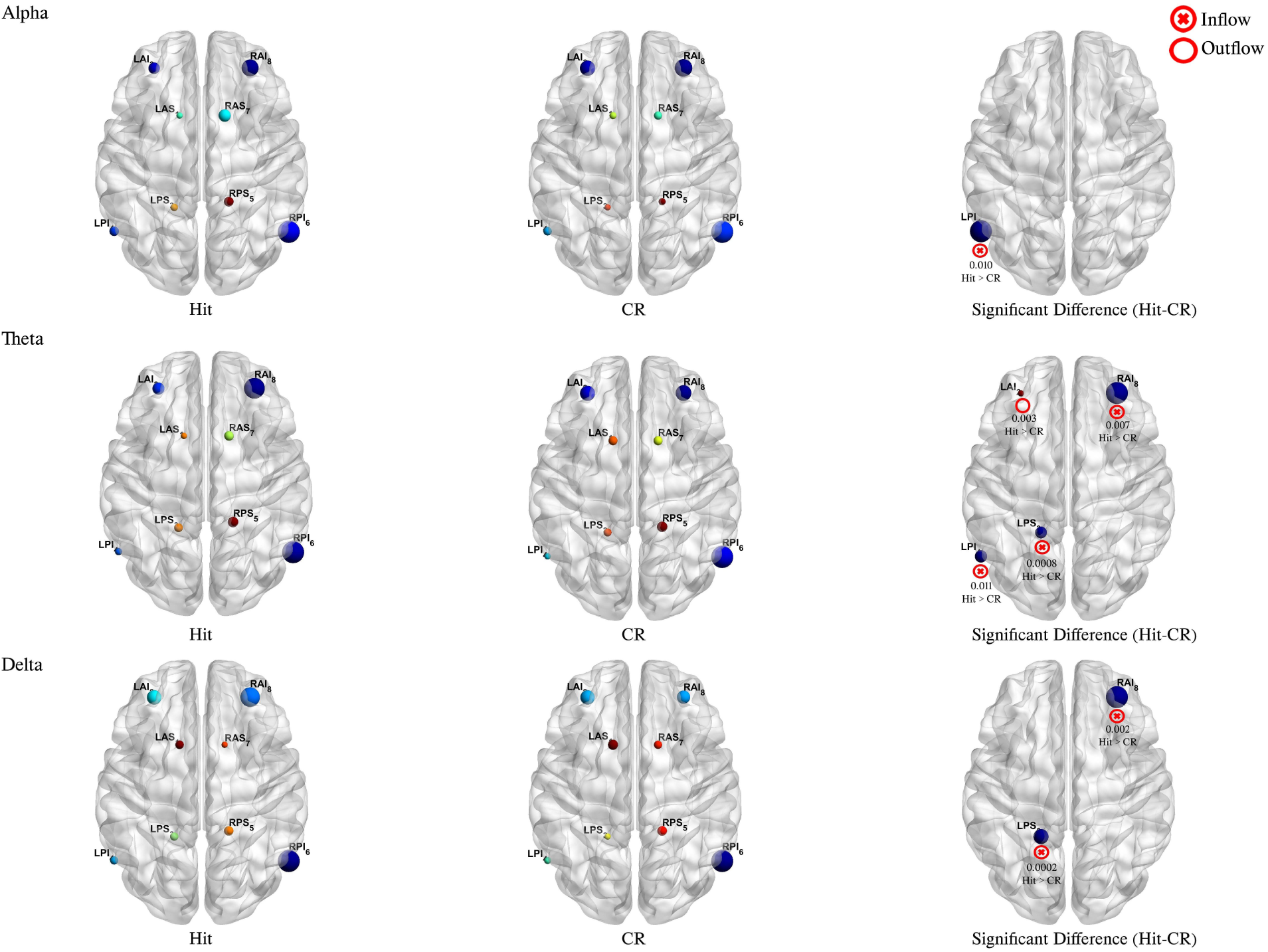
The information flow of regions estimated by dDTF in old/new (as Hit/CR) and the significant nodes between two conditions. The color of nodes present the outflow intensity and the node size shows inflow. Also the significant difference showed by special signs for inflow/outflow.

**Figure 6:**
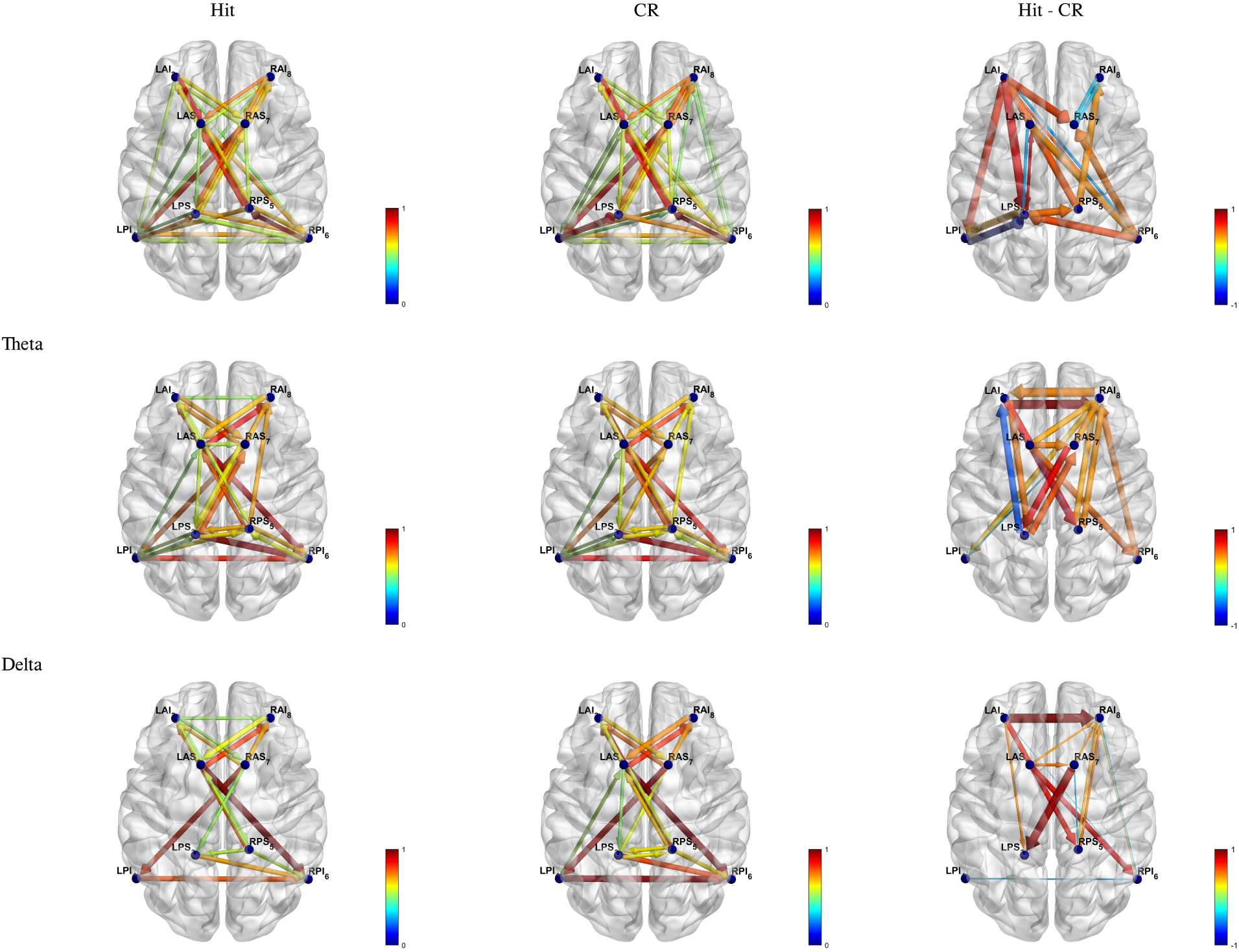
Average networks for old/new and their difference (*connectivity_old_* — *connectivity_new_*) by using the GPDC estimator, the difference is normalized for visualization, the color of edges present this difference and colors limited between blue (−1)/(0) and red (+1) so in third column red color means the old condition have a stronger connection than new and blue shows new have stronger connection at that edge.

**Figure 7:**
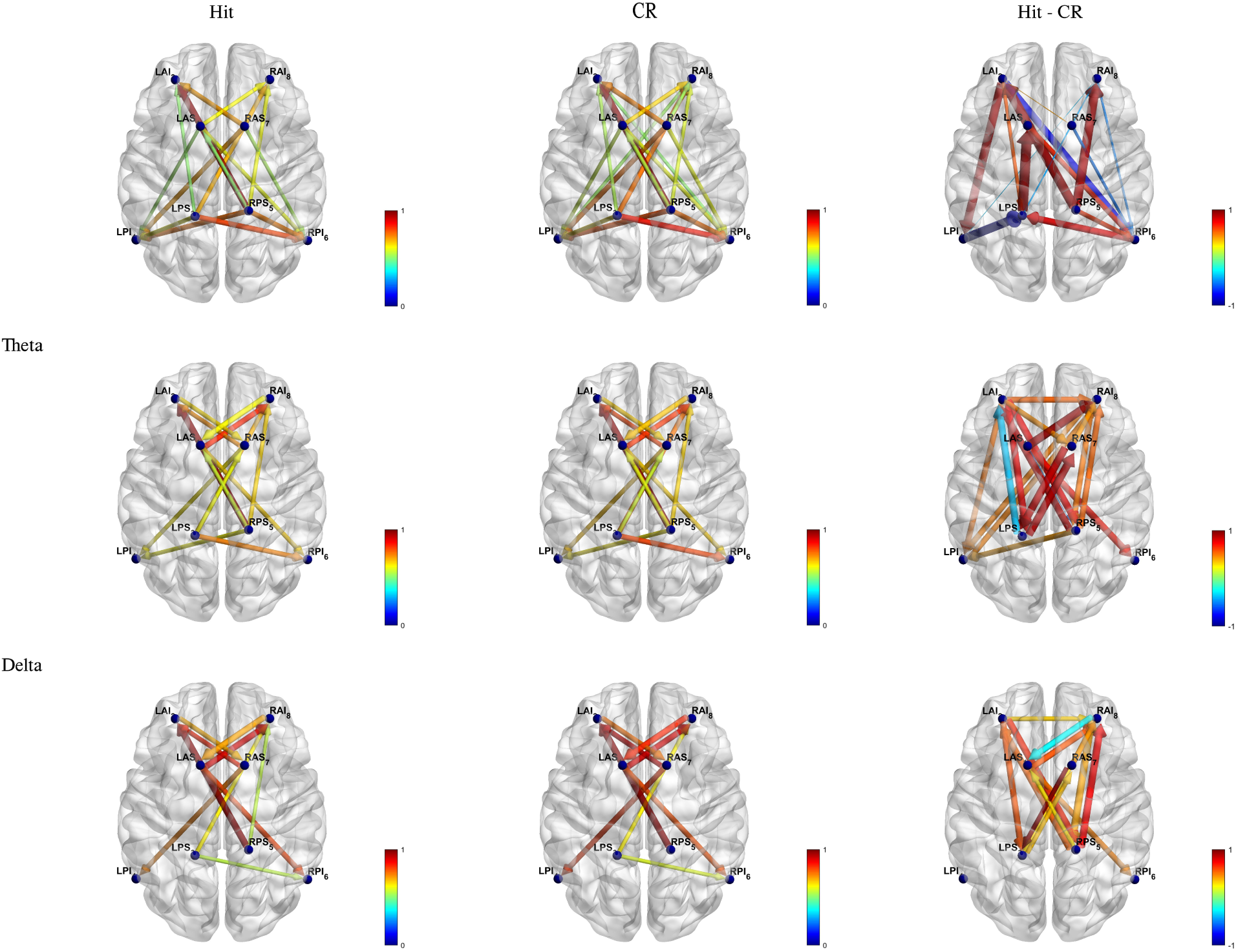
Average Networks for old/new and their difference (*connectivity_o1d_* — *connectivity_new_*) by using the dDTF estimator, the difference is normalized for visualization, the color of edges present this difference and colors limited between blue (−1)/(0) and red (+1) so in third column red color means the old condition have a stronger connection than new and blue shows new have stronger connection at that edge.

The GPDC network shows strong connection between Left and Right cortex. It shows strong connections between LAS to LPI in old/new and RAS to LPI.

The dDTF network shows strong connection between Left and Right cortex. Most strong connections are in anterior in low frequency bands and in posterior in high frequency band.

## 4 Discussion

In this study, we searched for determining the functional network interactions in brain regions previously involved in successful episodic memory retrieval and investigating how old/new word retrieval process have the desired results. Causal connectivity in frequency bands was estimated and compared between the two conditions. Evidence shows that EEG oscillations in specific frequency bands are related to memory process [46, 36, 32], Here memory process from the view of brain regional connection in frequency bands is investigated across twenty subjects and important results are obtained and reported.

The Theta and Gamma band are suggested as the pathways for brain region to interact [46, 43] for encoding and reveal memory, in our result, most of the significant connections are in these two bands. Our finding by GDDC and dDTF confirm the result from [15] by using Phase Lag Index as another connectivity estimator, they provide evidence that memory is highly correlated to the theta frequency band.

In our data both old/new are correct retrieval of memory, the old state involves recollection information while the new state is based on familiarity [16]. It is shown that recollection has stronger functional connectivity in gamma band between frontal and parietal cortex [13, 2], their task was very similar to our data and our result confirms this finding plus that shows the direction of connection between brain areas and the estimators have solved the problem of bivariate estimators.

The left frontal area has special role in learning and memory [13, 24, 27], especially in theta frequency band [40] and between prefrontal and temporal lobe [5], the dDTF result clearly shows the role of Left prefrontal cortex and its connection to parietal and temporal lobe.

The superior frontal gyrus is a hub for episodic memory-related network [38]. In another brain connectivity study for correct/incorrect episodic memory task, medial temporal lobe act as a hub for retrieval network [58]. In our result, especially the dDTF results show that the LAI node has a special role in the old/new difference.

In Semantic memory retrieval, it is shown low-frequency brain network works in the frontal cortex and high-frequency network active in all cortex [31]. The dDTF estimator shows very active frontal nodes in low frequency between frontal and pre-frontal to parietal regions and high-frequency network involve pre-frontal parietal and temporal cortex.

In [58, 38] they show stronger global connectivity for successful retrieval, these results are consistent with the studies about connectivity in episodic memory as well, because We observed the almost increase in the connections of the theta and gamma frequency bands. In another study [30], they reported that in the encoding phase of patients with Alzheimer’s, there are more connections in the delta, theta and alpha bands and they have strong connection in the retrieval phase in the Delta and Theta bands, but in the retrieval phase of the old words, our results show a stronger connection in the Delta and Theta bands. New words that are correctly rejected show a stronger connection in the gamma frequency bands in left parietal cortex, again like the old one. maybe in that condition subjects try to use some compensation mechanism to remember the items [30].

The information flows of region are local features and show which node have more input from or output of other regions. In our result GPDC inflow shows that the delta and theta band in RAI and LPS have special role in old/new difference, and other side dDTF outflow show the LAI in theta band has special node for recognition memory. It is inline with other research [60, 27, 40, 12, 50]. in this paper we shows more information that how they connect and work with other regions in detail.

To the best of our knowledge and exacting literature, there is a divergence of result and opinion in this field, these may become from the difference between tasks or connectivity estimator, to address the task issue we have used a reliable data that most of the psychological effects are considered and for connectivity estimator we have used two multivariate directed robust estimator that considered most of the issues in bivariate and undirected methods. we also should mention that the EEG signals not necessarily show accurate brain signals and our method maybe suffer from volume conducting.

## 5 Conclusion

Recent studies on functional connectivity have shown very valuable results but still, a lack of reliable results is felt because of little evidence and diversity in the results, to address these issue we conducted multivariate effective connectivity study to estimate the interaction between brain regions in the memory task. we found that theta and gamma band have a crucial role in old/new conditions. In the theta band, cross-connections are more activated in the old state and left frontal cortex acts as a hub for this system, the connections in this band indicating that in the old state the two hemispheres are interconnected, this finding is less discussed in the previous studies.

We found that old word memory retrieval was characterized by increases in network effective connectivity than new words in the delta, theta, gamma bands and decrease in the gamma band in the left temporal-parietal connection. the connections in theta band are between frontal to parietal cortex while the connections in the gamma band widespread in the left cortex.

Overall, our findings of change network connectivity during old vs. new word retrieval support the idea of global–specific changes in the memory task.

## 7 Author Contributions

T.C. and A.M.N. conceived of the presented idea. T.C. developed the theory and A.M.N Recommend the computations. M.J.D. verified and performed the analytical methods and prepare manuscript. T.C. and A.M.N. supervised the findings of this work. All authors discussed the results and contributed to the final manuscript.

## References

[1] Laura Astolfi et al. “Comparison of different cortical connectivity estimators for high-resolution EEG recordings”. In: Human Brain Mapping 28.2 (Feb. 2007), pp. 143–157. ISSN: 10659471. DOI: 10.1002/hbm.20263. URL: http://doi.wiley.com/10.1002/hbm.20263.

[2] Claudio Babiloni et al. “Functional frontoparietal connectivity during short-term memory as revealed by high-resolution EEG coherence analysis.” In: Behavioral neuroscience 118.4 (2004), p. 687.

[3] Luiz A. Baccala, K. Sameshima, and D. Y. Takahashi. “Generalized Partial Directed Coherence”. In: 2007 15th International Conference on Digital Signal Processing. IEEE, July 2007, pp. 163–166. ISBN: 1-4244-0881-4. DOI: 10.1109/ICDSP.2007.4288544. URL:http://ieeexplore.ieee.org/document/4288544/.

[4] Luiz A. Baccalá and Koichi Sameshima. “Partial directed coherence: a new concept in neural structure determination”. In: Biological Cybernetics 84.6 (May 2001), pp. 463–474. ISSN: 0340-1200. DOI: 10.1007/PL00007990. URL: http://link.springer.com/10.1007/PL00007990.

[5] Alexander R Backus et al. “Hippocampal-prefrontal theta oscillations support memory integration”. In: Current Biology 26.4 (2016), pp. 450–457.

[6] Lionel Barnett and Anil K. Seth. “Behaviour of Granger causality under filtering: Theoretical invariance and practical application”. In: Journal of Neuroscience Methods 201.2 (Oct. 2011), pp. 404–419. ISSN: 0165-0270. DOI: 10.1016/J.JNEUMETH.2011.08.010. URL: https://www.sciencedirect.com/science/article/pii/S0165027011004687.

[7] Erol Basar et al. “Gamma, alpha, delta, and theta oscillations govern cognitive processes”. In: International Journal of Psychophysiology 39.2-3 (Jan. 2001), pp. 241–248. ISSN: 0167-8760. DOI: 10.1016/S0167-8760(00)00145-8. URL: https://www.sciencedirect.com/science/article/pii/S0167876000001458.

[8] Yoav Benjamini and Yosef Hochberg. “Controlling the False Discovery Rate: A Practical and Powerful Approach to Multiple Testing”. In: Journal of the Royal Statistical Society: Series B (Methodological) 57.1 (Jan. 1995), pp. 289–300. ISSN: 00359246. DOI: 10.1111/j.2517-6161.1995.tb02031.x. URL: http://doi.wiley.com/10.1111/j.2517-6161.1995.tb02031.x.

[9] Carolyn JM Best et al. “Molecular alterations in primary prostate cancer after androgen ablation therapy”. In: Clinical Cancer Research 11.19 (2005), pp. 6823–6834.

[10] Nima Bigdely-Shamlo et al. “The PREP pipeline: standardized preprocessing for large-scale EEG analysis”. In: Frontiers in Neuroinformatics 9 (June 2015), p. 16. ISSN: 1662-5196. DOI: 10.3389/fninf.2015.00016. URL: http://journal.frontiersin.org/Article/10.3389/fninf.2015.00016/abstract.

[11] Katarzyna J Blinowska and Maciej Kamiński. “15 Multivariate Signal Analysis by Parametric Models”. In: Handbook of Time Series Analysis: Recent Theoretical Developments and Applications (2006), p. 373.

[12] Robert S. Blumenfeld and Charan Ranganath. “Prefrontal Cortex and Long-Term Memory Encoding: An Integrative Review of Findings from Neuropsychology and Neuroimaging”. In: The Neuroscientist 13.3 (June 2007), pp. 280–291. ISSN: 1073-8584. DOI: 10.1177/1073858407299290. URL: http://journals.sagepub.com/doi/10.1177/1073858407299290.

[13] Adrian P Burgess and Lia Ali. “Functional connectivity of gamma EEG activity is modulated at low frequency during conscious recollection”. In: International Journal of Psychophysiology 46.2 (2002), pp. 91–100.

[14] György Buzsáki. “The hippocampo-neocortical dialogue”. In: Cerebral cortex 6.2 (1996), pp. 81–92.

[15] Menorca Chaturvedi et al. “Phase Lag Index and Spectral power as QEEG features for identification of patients with Mild Cognitive Impairment in Parkinsońs disease”. In: Clinical Neurophysiology (2019).

[16] Tim Curran et al. “Combined pharmacological and electrophysiological dissociation of familiarity and recollection.” In: The Journal of neuroscience: the official journal of the Society for Neuroscience 26.7 (Feb. 2006), pp. 1979–85. ISSN: 1529-2401. DOI: 10.1523/JNEUROSCI.5370-05.2006. URL: http://www.ncbi.nlm.nih.gov/pubmed/16481430.

[17] M.J. Darvishi, A.M. Nasrabadi, and T. Curran. “Effective connectivity measuring of ERP signals in recognition memory process by Generalized Partial Directed Coherence”. In: 2016 23rd Iranian Conference on Biomedical Engineering and 20161 st International Iranian Conference on Biomedical Engineering (ICBME). IEEE, 2016, pp. 64–68. ISBN: 978-1-5090-3452-9. DOI: 10.1109/ICBME.2016.7890930. URL: http://ieeexplore.ieee.org/document/7890930/.

[18] Arnaud Delorme and Scott Makeig. “EEGLAB: an open source toolbox for analysis of single-trial EEG dynamics including independent component analysis”. In: Journal of Neuroscience Methods 134.1 (Mar. 2004), pp. 9–21. ISSN: 0165-0270. DOI: 10.1016/J.JNEUMETH.2003.10.009. URL: https://www.sciencedirect.com/science/article/pii/S0165027003003479.

[19] Arnaud Delorme et al. “EEGLAB, SIFT, NFT, BCILAB, and ERICA: New Tools for Advanced EEG Processing”. In: Computational Intelligence and Neuroscience 2011 (2011), pp. 1–12. ISSN: 1687-5265. DOI: 10.1155/2011/130714. URL:http://www.hindawi.com/journals/cin/2011/130714/.

[20] Audrey Duarte, Charan Ranganath, and Robert T. Knight. “What Neural Correlates Underlie Successful Encoding and Retrieval? A Functional Magnetic Resonance Imaging Study Using a Divided Attention Paradigm”. In: Journal of Neuroscience 23.6 (Sept. 2005), pp. 2407–2415. ISSN: 0270-6474. DOI: 10.1523/jneurosci.1392-05.2005. URL: https://www.jneurosci.org/content/25/36/8333.short.

[21] H. Eichenbaum, A.P. Yonelinas, and C. Ranganath. “The Medial Temporal Lobe and Recognition Memory”. In: Annual Review of Neuroscience 30.1 (July 2007), pp. 123–152. ISSN: 0147-006X. DOI: 10.1146/annurev.neuro.30.051606.094328. URL: http://www.annualreviews.org/doi/10.1146/annurev.neuro.30.051606.094328.

[22] Howard Eichenbaum. “A cortical-hippocampal system for declarative memory”. In: Nature Reviews Neuroscience 1.1 (Oct. 2000), pp. 41–50. ISSN: 1471-003X. DOI: 10.1038/35036213. URL: http://www.nature.com/articles/35036213.

[23] E E Fanselow et al. “Thalamic bursting in rats during different awake behavioral states.” In: Proceedings of the National Academy of Sciences of the United States of America 98.26 (Dec. 2001), pp. 15330–5. ISSN: 0027-8424. DOI: 10.1073/pnas.261273898. URL: http://www.ncbi.nlm.nih.gov/pubmed/11752471‥20http://www.pubmedcentral.nih.gov/articlerender.fcgi?artid=PMC65029.

[24] Emily A Farris et al. “Functional connectivity between the left and right inferior frontal lobes in a small sample of children with and without reading difficulties”. In: Neurocase 17.5 (2011), pp. 425–439.

[25] Juergen Fell and Nikolai Axmacher. “The role of phase synchronization in memory processes”. In: Nature Reviews Neuroscience 12.2 (Feb. 2011), pp. 105–118. ISSN: 1471-003X. DOI: 10.1038/nrn2979. URL: http://www.nature.com/articles/nrn2979.

[26] Esther Florin et al. “The effect of filtering on Granger causality based multivariate causality measures”. In: NeuroImage 50.2 (Apr. 2010), pp. 577–588. ISSN: 1053-8119. DOI: 10.1016/J.NEUROIMAGE.2009.12.050. URL: https://www.sciencedirect.com/science/article/pii/S1053811909013391.

[27] Nicolai Franzmeier et al. “Left frontal hub connectivity during memory performance supports reserve in aging and mild cognitive impairment”. In: Journal of Alzheimer’s Disease 59.4 (2017), pp. 1381–1392.

[28] C. W. J. Granger. “Investigating Causal Relations by Econometric Models and Cross-spectral Methods”. In: Econometrica 37.3 (Aug. 1969), p. 424. ISSN: 00129682. DOI: 10.2307/1912791. URL: https://www.jstor.org/stable/1912791?origin=crossref.

[29] Clive WJ Granger. “Testing for causality: a personal viewpoint”. In: Journal of Economic Dynamics and control 2 (1980), pp. 329–352.

[30] Yuliang Han et al. “Changes of EEG Spectra and Functional Connectivity during an Object-Location Memory Task in Alzheimer’s Disease”. In: Frontiers in Behavioral Neuroscience 11 (May 2017), p. 107. ISSN: 1662-5153. DOI: 10.3389/fnbeh.2017.00107. URL: http://journal.frontiersin.org/article/10.3389/fnbeh.2017.00107/full.

[31] Samer Hanouneh et al. “EEG power and functional connectivity correlates with semantic long-term memory retrieval”. In: Ieee Access 6 (2018), pp. 8695–8703.

[32] Gregor M. Hoerzer et al. “Directed coupling in local field potentials of macaque V4 during visual short-term memory revealed by multivariate autoregressive models”. In: Frontiers in Computational Neuroscience 4 (May 2010), p. 14. ISSN: 16625188. DOI: 10.3389/fncom.2010.00014. URL: http://journal.frontiersin.org/article/10.3389/fncom.2010.00014/abstract.

[33] J Benjamin Hutchinson, Melina R Uncapher, and Anthony D Wagner. “Posterior parietal cortex and episodic retrieval: convergent and divergent effects of attention and memory.” In: Learning & memory (Cold Spring Harbor, N.Y.) 16.6 (June 2009), pp. 343–56. ISSN: 1549-5485. DOI: 10.1101/lm.919109. URL: http://www.ncbi.nlm.nih.gov/pubmed/19470649%20http://www.pubmedcentral.nih.gov/articlerender.fcgi?artid=PMC2704099.

[34] M. J. Kaminski and K. J. Blinowska. “A new method of the description of the information flow in the brain structures”. In: Biological Cybernetics 65.3 (July 1991), pp. 203–210. ISSN: 0340-1200. DOI: 10.1007/BF00198091. URL:http://link.springer.com/10.1007/BF00198091.

[35] Maciej Kaminski and Katarzyna J. Blinowska. “The Influence of Volume Conduction on DTF Estimate and the Problem of Its Mitigation”. In: Frontiers in Computational Neuroscience 11 (May 2017), p. 36. ISSN: 1662-5188. DOI: 10.3389/fncom.2017.00036. URL: http://journal.frontiersin.org/article/10.3389/fncom.2017.00036/full.

[36] Wolfgang Klimesch. “EEG alpha and theta oscillations reflect cognitive and memory performance: a review and analysis”. In: Brain Research Reviews 29.2-3 (Apr. 1999), pp. 169–195. ISSN: 0165-0173. DOI: 10.1016/S0165-0173(98)00056-3. URL: https://www.sciencedirect.com/science/article/pii/S0165017398000563.

[37] Anna Korzeniewska et al. “Determination of information flow direction among brain structures by a modified directed transfer function (dDTF) method”. In: Journal of neuroscience methods 125.1-2 (2003), pp. 195–207.

[38] Chung-Yeon Lee and Byoung-Tak Zhang. “Effective EEG connectivity analysis of episodic memory retrieval”. In: Proceedings of the Annual Meeting of the Cognitive Science Society. Vol. 36. 36. 2014.

[39] John H McDonald. Handbook of biological statistics. Vol. 2. sparky house publishing Baltimore, MD, 2009.

[40] Lars Meyer et al. “Frontal–posterior theta oscillations reflect memory retrieval during sentence comprehension”. In: Cortex 71 (2015), pp. 205–218.

[41] Karen J. Mitchell and Marcia K. Johnson. “Source monitoring 15 years later: What have we learned from fMRI about the neural mechanisms of source memory?” In: Psychological Bulletin 135.4 (2009), pp. 638–677. ISSN: 1939-1455. DOI: 10.1037/a0015849. URL:http://doi.apa.org/getdoi.cfm?doi=10.1037/a0015849.

[42] Partha Mitra. Observed brain dynamics. Oxford University Press, 2007.

[43] Morris Moscovitch et al. “Episodic memory and beyond: the hippocampus and neocortex in transformation”. In: Annual review of psychology 67 (2016), pp. 105–134.

[44] Tim R. Mullen et al. “Real-time neuroimaging and cognitive monitoring using wearable dry EEG”. In: IEEE Transactions on Biomedical Engineering 62.11 (Nov. 2015), pp. 2553–2567. ISSN: 0018-9294. DOI: 10.1109/TBME.2015.2481482. URL:http://ieeexplore.ieee.org/document/7274673/.

[45] Kenneth A. Norman and Randall C. O’Reilly. “Modeling hippocampal and neocortical contributions to recognition memory: A complementary-learning-systems approach.” In: Psychological Review 110.4 (2003), pp. 611–646. ISSN: 1939-1471. DOI: 10.1037/0033-295X.110.4.611. URL: http://doi.apa.org/getdoi.cfm?doi=10.1037/0033-295X.110.4.611.

[46] Erika Nyhus and Tim Curran. “Functional role of gamma and theta oscillations in episodic memory”. In: Neuroscience & Biobehavioral Reviews 34.7 (2010), pp. 1023–1035.

[47] Hae-Jeong Park and Karl Friston. “Structural and functional brain networks: from connections to cognition.” In: Science (New York, N.Y.) 342.6158 (Nov. 2013), p. 1238411. ISSN: 1095-9203. DOI: 10.1126/science.1238411. URL: http://www.ncbi.nlm.nih.gov/pubmed/24179229.

[48] Guillaume A Rousselet. “Does filtering preclude us from studying ERP time-courses?” In: Frontiers in psychology 3 (2012), p. 131.

[49] Jon S. Simons and Hugo J. Spiers. “Prefrontal and medial temporal lobe interactions in long-term memory”. In: Nature Reviews Neuroscience 4.8 (Aug. 2003), pp. 637–648. ISSN: 1471-003X. DOI: 10.1038/nrn1178. URL: http://www.nature.com/articles/nrn1178.

[50] Julia Spaniol et al. “Event-related fMRI studies of episodic encoding and retrieval: Meta-analyses using activation likelihood estimation”. In: Neuropsychologia 47.8-9 (July 2009), pp. 1765–1779. ISSN: 0028-3932. DOI: 10.1016/J.NEUROPSYCHOLOGIA.2009.02.028. URL: https://www.sciencedirect.com/science/article/pii/S0028393209001067.

[51] Michael C. Stevens. “The developmental cognitive neuroscience of functional connectivity”. In: Brain and Cognition 70.1 (June 2009), pp. 1–12. ISSN: 0278-2626. DOI: 10.1016/J.BANDC.2008.12.009. URL: https://www.sciencedirect.com/science/article/pii/S0278262608003369.

[52] Nasibeh Talebi, Ali Motie Nasrabadi, and Iman Mohammad-Rezazadeh. “Estimation of effective connectivity using multi-layer perceptron artificial neural network”. In: Cognitive Neurodynamics 12.1 (Feb. 2018), pp. 21–42. ISSN: 1871-4080. DOI: 10.1007/s11571-017-9453-1. URL: http://link.springer.com/10.1007/s11571-017-9453-1.

[53] Darren Tanner, Kara Morgan-Short, and Steven J. Luck. “How inappropriate high-pass filters can produce artifactual effects and incorrect conclusions in ERP studies of language and cognition”. In: Psychophysiology 52.8 (Aug. 2015), pp. 997–1009. ISSN: 00485772. DOI: 10.1111/psyp.12437. URL: http://doi.wiley.com/10.1111/psyp.12437.

[54] James Theiler et al. “Testing for nonlinearity in time series: the method of surrogate data”. In: Physica D: Nonlinear Phenomena 58.1-4 (Sept. 1992), pp. 77–94. ISSN: 0167-2789. DOI: 10.1016/0167-2789(92)90102-S. URL: https://www.sciencedirect.com/science/article/pii/016727899290102S.

[55] Rufin VanRullen. “Four Common Conceptual Fallacies in Mapping the Time Course of Recognition”. In: Frontiers in Psychology 2 (Dec. 2011), p. 365. ISSN: 1664-1078. DOI: 10.3389/fpsyg.2011.00365. URL: http://journal.frontiersin.org/article/10.3389/fpsyg.2011.00365/abstract.

[56] Kaia L. Vilberg and Michael D. Rugg. “Memory retrieval and the parietal cortex: A review of evidence from a dual-process perspective”. In: Neuropsychologia 46.7 (June 2008), pp. 1787–1799. ISSN: 0028-3932. DOI: 10.1016/J.NEUROPSYCHOLOGIA.2008.01.004. URL: https://www.sciencedirect.com/science/article/pii/S0028393208000158.

[57] Ioannis Vlachos et al. “The concept of effective inflow: application to interictal localization of the epileptogenic focus from iEEG”. In: IEEE Transactions on Biomedical Engineering 64.9 (2016), pp. 2241–2252.

[58] Andrew J Watrous et al. “Frequency-specific network connectivity increases underlie accurate spatiotemporal memory retrieval”. In: Nature Neuroscience 16.3 (Mar. 2013), pp. 349–356. ISSN: 1097-6256. DOI: 10.1038/nn.3315. URL: http://www.nature.com/articles/nn.3315.

[59] Mark E Wheeler and Randy L Buckner. “Functional-anatomic correlates of remembering and knowing”. In: NeuroImage 21.4 (Apr. 2004), pp. 1337–1349. ISSN: 1053-8119. DOI: 10.1016/J.NEUROIMAGE.2003.11.001. URL: https://www.sciencedirect.com/science/article/pii/S1053811903007213.

[60] Mark E Wheeler et al. “Functional dissociation among components of remembering: control, perceived oldness, and content.” In: The Journal of neuroscience: the official journal of the Society for Neuroscience 23.9 (May 2003), pp. 3869–80. ISSN: 1529-2401. DOI: 10.1523/jneurosci.5295-04.2005. URL: http://www.ncbi.nlm.nih.gov/pubmed/12736357.

[61] Andreas Widmann, Erich Schröger, and Burkhard Maess. “Digital filter design for electrophysiological data – a practical approach”. In: Journal of Neuroscience Methods 250 (July 2015), pp. 34–46. ISSN: 0165-0270. DOI: 10.1016/J.JNEUMETH.2014.08.002. URL: https://www.sciencedirect.com/science/article/pii/S0165027014002866.

[62] Tal Yarkoni et al. “Cognitive neuroscience 2.0: building a cumulative science of human brain function”. In: Trends in Cognitive Sciences 14.11 (Nov. 2010), pp. 489–496. ISSN: 1364-6613. DOI: 10.1016/J.TICS.2010.08.004. URL: https://www.sciencedirect.com/science/article/pii/S1364661310002019.

[63] Andrew P Yonelinas et al. “Separating the brain regions involved in recollection and familiarity in recognition memory”. In: Journal of Neuroscience 25.11 (2005), pp. 3002–3008.

